# MicroRNA-mRNA networks are dysregulated in opioid use disorder postmortem brain: further evidence for opioid-induced neurovascular alterations

**DOI:** 10.1101/2022.07.19.500679

**Authors:** Sandra L. Grimm, Emily F. Mendez, Laura Stertz, Thomas D. Meyer, Gabriel R. Fries, Tanmay Gandhi, Rupa Kanchi, Sudhakar Selvaraj, Antonio L. Teixeira, Tom Kosten, Preethi Gunaratne, Cristian Coarfa, Consuelo Walss-Bass

## Abstract

To understand mechanisms and identify potential targets for intervention in the current crisis of opioid use disorder (OUD), postmortem brains represent an under-utilized resource. To refine previously reported gene signatures of neurobiological alterations in OUD from the dorsolateral prefrontal cortex (Brodmann Area 9, BA9), we explored the role of microRNAs (miRNA) as powerful epigenetic regulators of gene function. Building on the growing appreciation that miRNAs can cross the blood-brain barrier, we carried out miRNA profiling in same-subject postmortem samples from BA9 and blood tissues. miRNA-mRNA network analysis showed that even though miRNAs identified in BA9 and blood were fairly distinct, their target genes and corresponding enriched pathways were highly overlapping, with tube development and morphogenesis, and pathways related to endothelial cell function and vascular organization, among the dominant enriched biological processes. These findings point to robust, redundant, and systemic opioid-induced miRNA dysregulation with potential functional impact on transcriptomic changes. Further, using correlation network analysis we identified cell-type specific miRNA targets, specifically in astrocytes, neurons, and endothelial cells, associated with OUD transcriptomic dysregulation. Our refined miRNA-mRNA networks enabled identification of novel pharmaco-chemical interventions for OUD, particularly targeting the TGF beta-p38MAPK signaling pathway. Finally, leveraging a collection of control brain transcriptomes from the Genotype-Tissue Expression (GTEx) project, we identified correlation of OUD miRNA targets with TGF beta, hypoxia, angiogenesis, coagulation, immune system and inflammatory pathways. These findings support previous reports of neurovascular and immune system alterations as a consequence of opioid abuse.

## Introduction

Opioid use disorder (OUD) represents a major public health problem with alarming rates of overdoses and deaths. The neurobiological consequences of long-term opioid abuse are not well understood. MicroRNAs (miRNAs) are 17-22 nucleotide sequences that act as pleiotropic regulators of mRNAs through imperfect matching, mostly at the 3’ UTR of genes (4-6). miRNAs are key modulators of intercellular communication across the brain (7) across multiple species (8) and are dysregulated in psychiatric disorders such as schizophrenia and bipolar disorder (9). Opioid exposure significantly alters miRNA expression in many regions of the brain, but most notably in the prefrontal cortex and the striatum. Several miRNAs involved in regulation of synaptic plasticity are hypothesized to underlie drug addiction (10) and miRNAs have been shown to regulate µ-opioid receptor levels and modulate opioid tolerance (11, 12). These studies point towards miRNAs as critical short-term and long-term epigenetic modulators of opioid effects in the brain through regulation of gene expression. Importantly, as miRNAs can cross the blood/brain barrier (BBB) (13, 14), potentially via exosomes (15-17), assessment of differential miRNA expression in blood can prove helpful to identify functional biomarkers of dysregulation of target genes in brain regions affected by OUD.

Bioinformatics pipelines are one emerging area that can substantially improve the signal-to-noise ratio on efforts to extract critical drivers of signaling pathways that underlie OUD-related changes in the brain. These pipelines have been developed to identify biosignatures and new druggable targets from miRNA-regulated networks inferred using miRNA and mRNA profiles measured in the same specimens. This approach exceeds both the single gene discovery approach as well as the use of large gene signatures that, due to their diffuse biological impact, might otherwise prove intractable for elucidation of mechanisms and effective repurposing of pharmaco-chemical agents.

To explore the potential role of miRNAs as regulators of transcription in opioid-exposed brains, we employed a multi-pronged sequencing and analytical approach. RNA-Sequencing (RNA-Seq) was carried out in bulk tissue from the dorsolateral prefrontal cortex (DLPFC, Brodmann area 9, BA9) of individuals with OUD and controls, as we previously described (18). smallRNA-Seq to identify miRNAs was also performed in both BA9 and blood collected from the same donors. Dysregulated miRNA networks were inferred, and miRNA target genes were further examined using pathway enrichment, repurposable drug identification, and correlation-based network analysis. An overview of our analytical approach is outlined in Figure 1.

**Figure 1.**
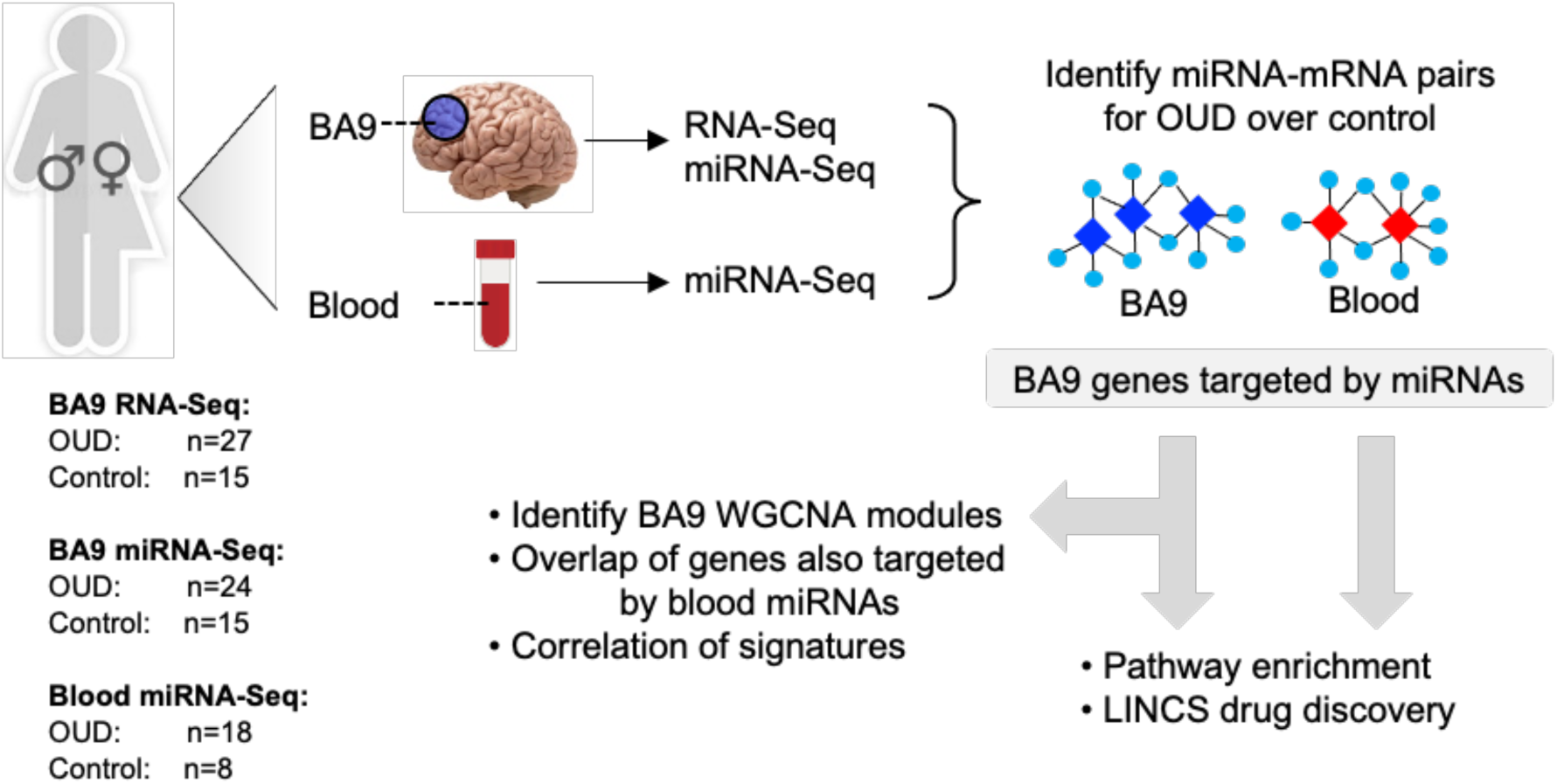
Analytical approach overview. RNA-Sequencing was carried out in brain tissue collected postmortem from individuals with OUD and controls. smallRNA-Seq was carried out in both brain and blood tissue from a subset of the same individuals. Dysregulated miRNA-RNA networks were inferred, and miRNA target genes were further investigated using pathway enrichment, repurposable drug identification, and correlation-based network analysis.

## Methods

### Sample information

Postmortem brain and peripheral blood tissues were obtained from the University of Texas Health Science Center at Houston (UTHealth) Brain Collection in collaboration with the Harris County Institute of Forensic Science, with consent from the next of kin (NOK) and approval from the Institutional Review Board. Medical Examiner Reports, including toxicology, were obtained and a detailed Psychological Autopsy was performed on the donor by interviewing the NOK (19). Psychiatric clinical phenotypes (evidence of depression, mania, psychosis), age of onset of drug use, types of drugs used, smoking and drinking history, and any co-morbidities were obtained. A diagnosis of OUD, or designation as non-psychiatric control, was confirmed by the psychological autopsy performed and a consensus meeting with three trained clinicians.

For this project, we focused on the dorsolateral prefrontal cortex (DLPFC, Brodmann area 9, BA9) because this is a key cerebral area involved in inhibition and impulsivity, functions that are altered in substance use disorders (20). Upon receipt of the brain, the right hemisphere was coronally sectioned, immediately frozen, and stored at -80° C. Dissections of BA9, defined within the DLPFC between the superior frontal gyrus and the cingulate sulcus, were obtained using a 4mm cortical punch, yielding approximately 100mg of tissue. Postmortem interval (PMI) was calculated from the estimated time of death until tissue preservation. Peripheral blood samples were collected into EDTA-containing tubes, and then stored as whole blood at -80 °C until further use.

### RNA-Seq analysis

We analyzed RNA-Seq data generated as previously described (Mendez et al 2021) for 15 controls and 27 OUD cases using bulk BA9 tissue. RNA-Seq data was trimmed for low quality base pairs and adapter sequences using trim_galore. Sequencing reads were mapped to the human genome build UCSC hg38 using STAR (21). Gene expression was quantified using featureCounts (22). The following demographic variables were accounted and regressed out using the R package RUVr (23): Gender, Age, PMI (hours), pH, and RNA integrity number (RIN). We used the R package EdgeR (24) to infer differentially expressed genes between groups, with significance achieved at a fold change exceeding 1.5x and FDR-adjusted p-value<0.2.

### smallRNA-Seq analysis

The small RNA fraction was isolated from bulk BA9 tissue from 15 controls and 24 OUD cases for which bulk RNA-Seq data was already available (18), and from blood from a subset of the same individuals (8 controls and 18 OUD). We then generated small RNA libraries using the Illumina small RNA protocol and sequenced the small RNA libraries on an Illumina Genome Analyzer NextSeq 500. We analyzed 3-4 million reads per sample using our published bioinformatics pipeline (25). We constructed RNA-Seq libraries using the Takara SMARTer Universal Low Input RNA Kit designed to handle 2–100 ng of total RNA and retain strand-specific information.

The smallRNA-Seq data was trimmed for low quality base pairs and adapter sequences using trim_galore. Sequencing reads were mapped to the human genome build UCSC hg38 using STAR (21) allowing for at most 10 matches. miRNA expression was quantified using featureCounts (22) and the miRBase reference database (5). The following demographic variables were accounted and regressed out using the R package RUVr (23): Gender, Age, PMI (hours), pH, and RNA integrity number (RIN). We used the R package EdgeR (24) to infer differentially expressed genes between groups. We screened for miRNA with p-value<0.05. We further conducted miRNA-mRNA integration using the SigTerms methodology (26), with significance achieved at FDR-adjusted p-value<0.05. miRNA-mRNA networks were visualized using the Cytoscape platform (27).

### Gene Network analysis

Gene network analysis was carried out using the Weighted Correlation Network Analysis (WGCNA) R package (28), using default parameters and a minimum module size of 5. Brain cell type composition was inferred with CIBERSORT (29) using reference gene signatures previously reported (30) for human brain cell types and default parameters. Significant association with clinical traits was performed using WGCNA, with significance achieved at p<0.05.

### Drug repurposing analysis

Repurposing of drugs targeting the gene signatures determined using the miRNA-mRNA analysis pipeline was performed using the Library of Integrated Network-Based Cellular Signatures (LINCS) Program/Connectivity Map platform (31).

### Signature correlation analysis

A methodology used frequently to assess the interaction between gene signatures in human cohorts is the correlation of gene signature scores; this approach has been utilized both in cancer (32-34) and non-cancer systems (35-37). We downloaded data for 425 control bulk BA9 brain tissues collected by the Genotype-Tissue Expression (GTEx) project (38), using the GTEx data portal. We computed signature scores separately for the 50 Hallmark pathways (39), then for the up and down gene targets of miRNAs and for WGCNA modules using summed z-scores. We then assessed the association using Pearson Correlation Coefficient, with significance achieved for p<0.05. Correlation heatmaps were plotted using the Python language scientific library.

## Results

Postmortem brain and blood tissues were obtained through the UT Health Brain Collection. Demographic information is provided in Table 1, with additional detailed information previously described (18). The controls were 87% male with an average age of 55 years, and the OUD group were 56% male with an average age of 39 years.

**Table 1.** Demographic information of samples used for RNA-Sequencing and smallRNA-Sequencing.

| **<u>Demographics</u>** | **<u>Control</u>** | **<u>OD</u>** |
| --- | --- | --- |
| Total number of samples | 15 | 27 |
| Males (%) | 13 (87%) | 15 (56%) |
| Age mean (sd) | 55 (14) | 39 (13) |
| PMI mean in hours (sd) | 29 (7) | 26 (9) |
| Ethnicity (White/Black/Hispanic/Asian) | 8/4/2/1 | 25/5/0/0 |

### miRNA-mRNA networks are dysregulated in OUD

We used RNA-sequencing to profile bulk BA9 tissue from 15 controls and 27 OUD cases; employing stringent criteria of FDR<0.2 and fold change exceeding 1.5x, we determined differential expression of 837 protein coding genes, with 144 induced in OUD and 693 decreased, after controlling for age, gender, PMI, RIN and pH (Figure 2, Supplemental Table 1A). Since our miRNA-mRNA integration network approach applied a stringent statistical criterion employing the Benjamini-Hochberg correction, we used a relaxed criterion of p<0.05 to screen for network candidates based on differential expression of miRNAs, controlling for the same demographic and clinical covariates. From this analysis, BA9-measured miRNAs yielded 89 differentially expressed miRNAs, with 29 induced in OUD and 60 suppressed (Figure 2C, Supplemental Table 1B), whereas blood-measured miRNAs yielded 104 differentially expressed miRNAs, with 51 increased in OUD and 53 decreased (Figure 2, Supplemental Table 1B). We used the SigTerms bioinformatics platform (25, 26), developed by our group, to extract biologically significant miRNAs and mRNA targets underlying OUD by integrating sequencing data with miRNA target prediction algorithms from mirDB (40) to extract miRNA-mRNA target pairs that are oppositely correlated in expression with the highest enrichment statistics. Using BA9 miRNA candidates (Figure 2A, C) we identified 7 OUD down-regulated miRNAs as mechanistic miRNAs that drove a network of 44 OUD up-regulated genes at FDR<0.05 (Figure 3A). Conversely, we identified 10 OUD up-regulated miRNAs that drove, at a stringent FDR<0.05, a network of 182 OUD down-regulated genes (Figure 3A). Using blood miRNA candidates (Figure 2A, D), we identified 5 OUD down-regulated miRNAs with targets enriched in 31 up-regulated coding genes and, conversely, we identified 21 OUD up-regulated miRNAs with targets enriched in 304 BA9 down-regulated genes (Figure 3B). We did not identify overlap between potential BA9 and blood miRNA regulators. However, overlap of their target genes was robust with 123 down-regulated genes and 12 up-regulated genes enriched for targets of both BA9 and blood miRNAs (Figure 3C).

**Figure 2.**
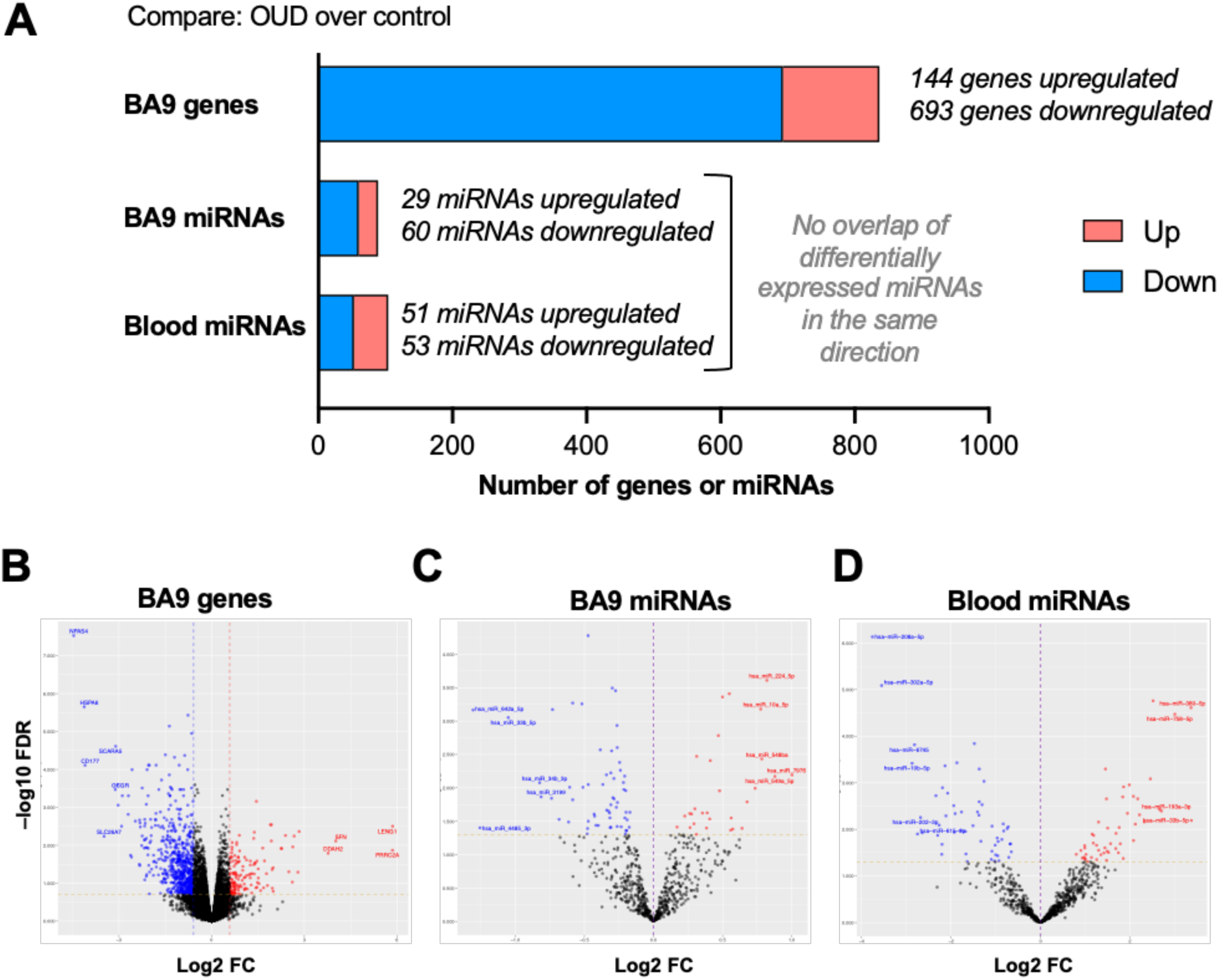
Differentially expressed protein-coding genes and microRNAs in brain and blood from individuals with OUD. (A) Summary of differentially expressed miRNA-mRNA network components. Volcano plots of BA9 protein coding genes (B), BA9 miRNAs (C), and blood miRNAs (D), comparing OUD to controls. The top 10 features by log2 fold change are labeled.

**Figure 3.**
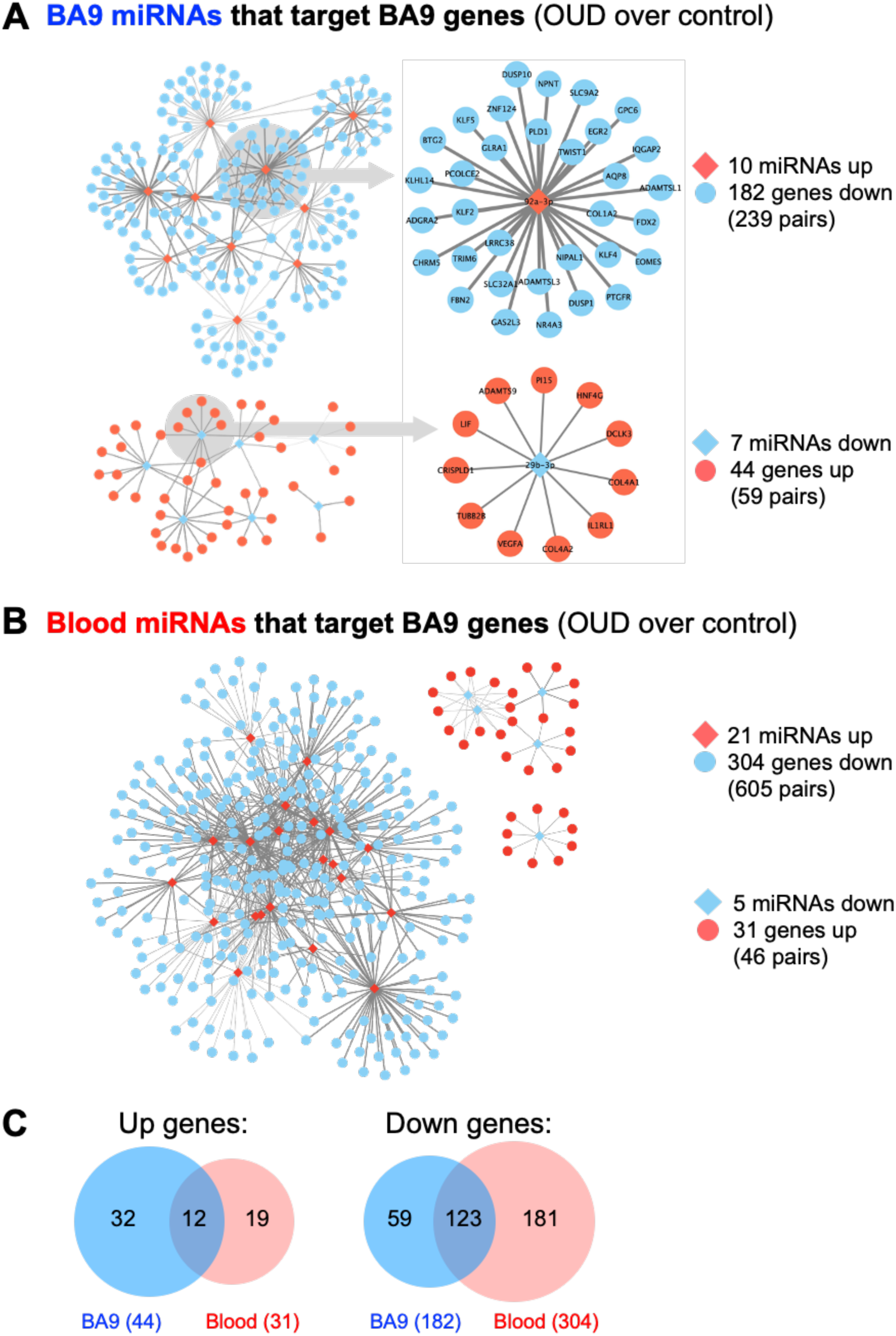
Dysregulated miRNA-mRNA networks in brains from OUD donors. Using the significant dysregulated protein coding genes measured in BA9, we determined enriched miRNA targets based on the OUD-associated miRNA candidates measured in (A) BA9 and (B) blood via the mirDB compendium. C. Overlap of up and down regulated genes targeted by miRNAs in BA9 and blood.

The direct (canonical) mode of miRNA-driven regulation of mRNA is that of transcription suppression, e.g., up-regulated miRNA targets are enriched in down-regulated genes and conversely down-regulated miRNA targets are enriched in up-regulated genes. However, indirect (non-canonical) regulation is possible via intermediate lincRNA modulation (41). As such, we also evaluated potential miRNA targets enrichment in coding genes changes in the same direction. Based on BA9 miRNA candidates, we identified 3 up-regulated miRNAs with targets enriched in 21 OUD up-regulated genes, and 27 down-regulated miRNAs with targets enriched in 329 down-regulated genes (Supplemental Figure 1A). Interestingly, the number of miRNA-mRNA pairs for non-canonical regulation exceeded the number for canonical regulation. Despite different modes of regulation, we further determined that 14 up-regulated genes and 134 down-regulated genes can be regulated both by canonical and non-canonical regulation of miRNAs (Supplemental Figure 1A, Supplemental Table 2A). Whereas a similar picture emerged using blood miRNA candidates, the difference between canonical and non-canonical miRNA-mRNA pairs was not as striking (Supplemental Figure 1B, Supplemental Table 2B). Specifically, 11 up-regulated miRNAs in blood targeted 62 OUD up-regulated genes, and 24 down-regulated miRNAs targeted 309 down-regulated genes. A redundancy between blood miRNA canonical and non-canonical gene targets was observed as well. Twenty-four OUD up-regulated genes and 206 OUD down-regulated genes could be targeted via both canonical and non-canonical miRNA-mRNA regulation.

### Pathway enrichment and drug repurposing via targeting miRNA-mRNA networks

Based on BA9 miRNAs/BA9 gene networks and blood miRNAs/BA9 gene networks, we determined enriched Gene Ontology pathways using the MSigDB approach based on hypergeometric distribution, with significance achieved at FDR<0.05 (Figure 4A, Supplemental Table 3). Interestingly, targets of both BA9 miRNAs and of blood miRNAs enriched for similar processes; whereas development and morphogenesis were the dominant dysregulated biological pathways and processes, other common terms among the top 25 enriched pathways were cell adhesion, lipid metabolism, and cell proliferation. Reassuringly, 1020 pathways were enriched in gene targets of both BA9 miRNAs and blood miRNAs (Figure 4B).

**Figure 4.**
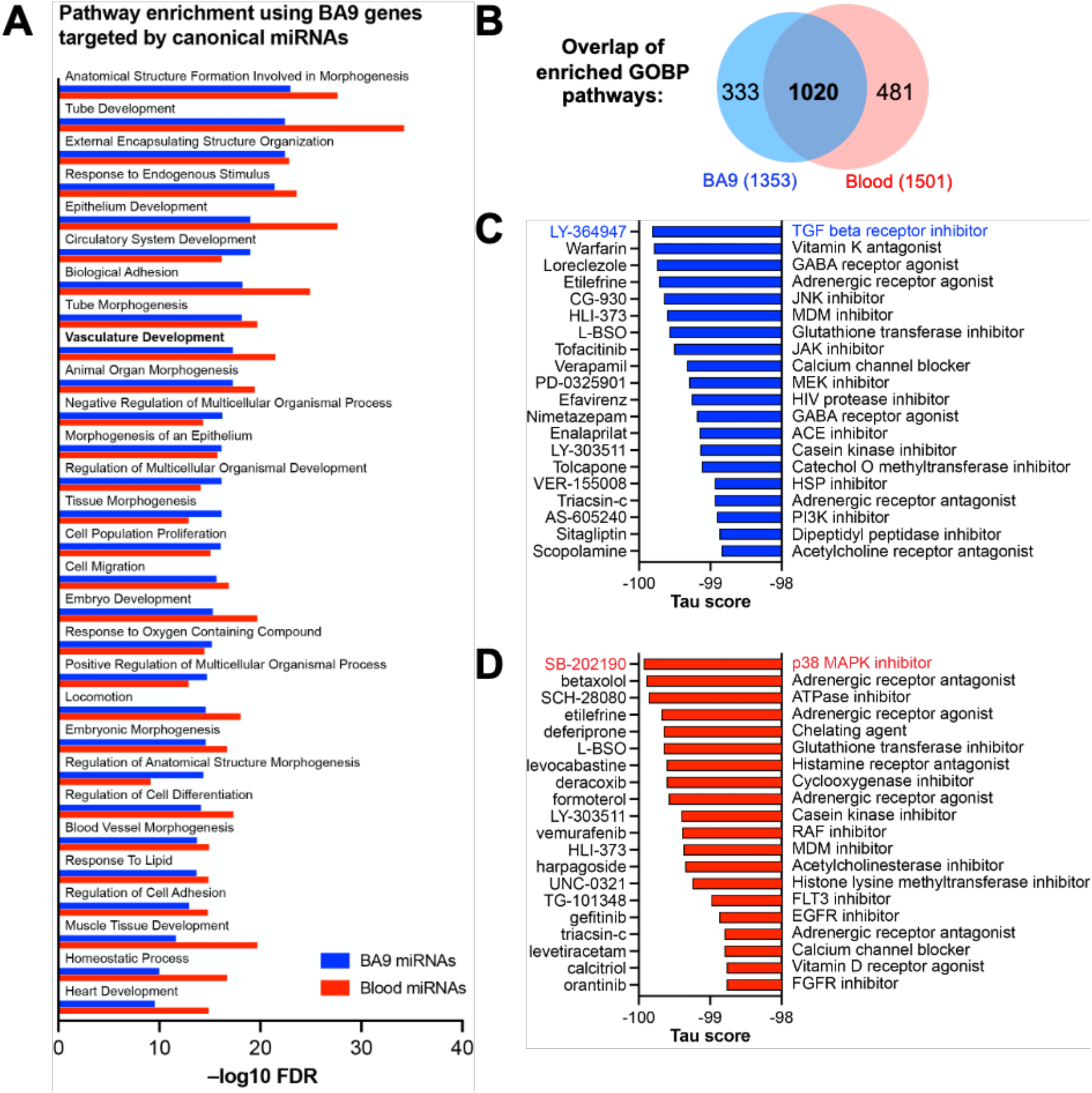
Integrative analysis of miRNA-mRNA networks. Pathway enrichment was carried out with the enriched gene targets of BA9 and blood miRNAs (A), using the Gene Ontology compendium, with overlap of enriched pathways shown in panel B. Repurposable drugs were ranked based on their anti-correlation with the gene targets detected in miRNA-mRNA networks from BA9 (C) and blood (D). mRNAs significant at FDR<0.2, and miRNAs significant at p<0.05. Networks significant at FDR<0.05.

Repurposing of drugs based on gene signatures using the Library of Integrated Network-Based Cellular Signatures (LINCS) L1000 database (31) has generated effective drug candidates in numerous disease systems, including alcohol use disorder (42, 43). Smaller, refined gene signatures could be a more effective to guide pharmaco-chemical interventions via drug repurposing. Thus, we interrogated the LINCS database with our miRNA-mRNA network genes generated based on either BA9 miRNAs (Figure 4C) or blood miRNAs (Figure 4D). The top drugs identified were a TGF beta receptor inhibitor for BA9 miRNAs, and a p38 MAPK inhibitor for blood miRNAs. Other classes of drugs were commonly found based on both networks, such as those targeting adrenergic receptors.

### WGCNA reveals miRNA-mRNA networks associated with individual cell types

Using CIBERSORT, we inferred cell type composition of the BA9 bulk tissue samples based on a collection of brain cell type signatures (30). Next, we employed Weighted Gene Correlation Network Analysis (WGCNA) (28) to identify modules in the BA9 miRNA-mRNA network, identifying seven modules ranging from 9 to 121 genes (Figure 5A). By using the brain cell type composition and the OUD status as traits, we performed module-trait association analysis, identifying WGCNA modules significantly associated with relative proportion of astrocytes, neurons, or endothelial cells and also with OUD status (Figure 5B). The yellow and green modules associated positively with astrocyte abundance and negatively with neuron abundance; overall the yellow, green, and blue modules associated positively with OUD status and comprised the majority of the canonical miRNA-mRNA network driven by down-regulated miRNAs and up-regulated genes (Figures 3A, 5C). The black module associated positively with neuron abundance and negatively with astrocyte abundance. The largest module, turquoise, and the red module associated positively with endothelial cells abundance. The turquoise, brown, red, and black modules associated negatively with OUD status and comprised the majority of the canonical miRNA-mRNA network driven by up-regulated miRNAs and down-regulated genes (Figure 3B, 5C).

**Figure 5.**
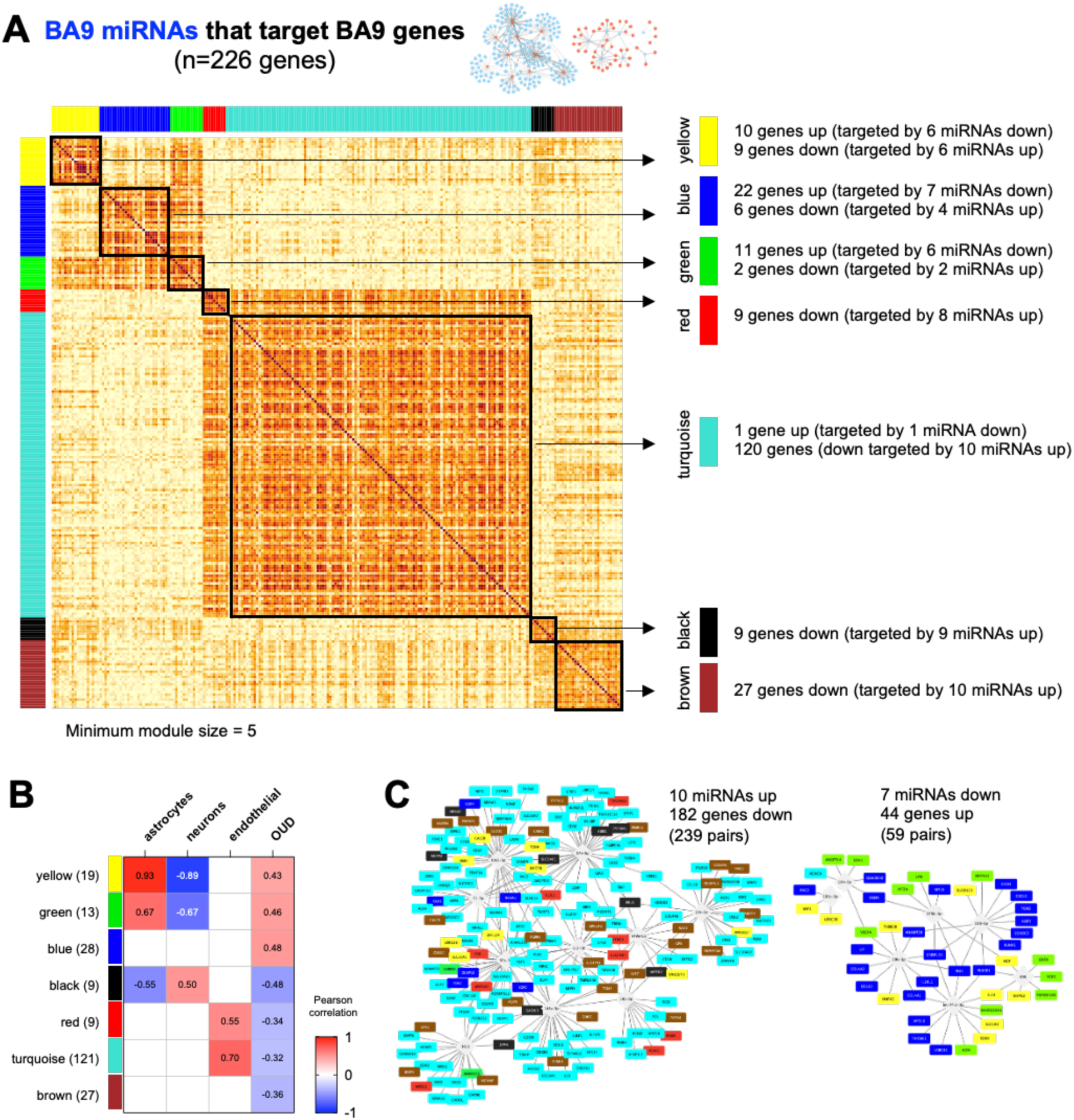
Correlation-based analysis of BA9 canonical miRNA-mRNA network genes. (A) Weighted gene correlation network analysis (WGCNA) was carried out for the 226 BA9 gene targets of BA9 miRNAs, identifying 7 distinct gene modules with a minimum module size of 5. (B) Association with brain cell type abundance and OUD diagnosis was evaluated for each module. (C) Networks comprised of increased miRNAs and decreased genes correlated with the turquoise, red, black, and brown modules. Networks comprised of decreased miRNAs and increased genes correlated with the yellow, blue, and green WGCNA modules.

As previously indicated, even though the differential miRNAs inferred from BA9 and blood were non-overlapping, 135 gene targets overlapped, with 123 down and 12 up (Figure 3C, Supplemental Figure 2). Robust overlaps were also observed between the BA9 WGCNA gene modules and blood miRNA-mRNA network targets; the black module was fully targeted by blood miRNAs (9/9 genes), and the turquoise module showed a 72% overlap (87/121 genes). The rest of the modules showed 31-56% overlap with blood miRNA gene targets (Supplemental Figure 2, Supplemental Table 4).

To assess if the WGCNA modules associated with distinct pathways, we first downloaded a database of 425 transcriptomes of control BA9 tissues compiled by the Genotype Tissue Expression (GTEx) project (38), then computed gene signature scores for OUD up-regulated and OUD down-regulated miRNA targets, respectively, for each of the WGCNA modules, and for a reference collection of the 50 Hallmark pathways (39). Strikingly, the signature scores for miRNA targets showed distinct correlation patterns based on directionality: OUD down-regulated miRNA targets correlated with hypoxia, TGF beta signaling, angiogenesis, and coagulation, whereas OUD up-regulated miRNA targets correlated with immune system and inflammatory pathways (Figure 6, Supplemental Figure 3).

**Figure 6.**
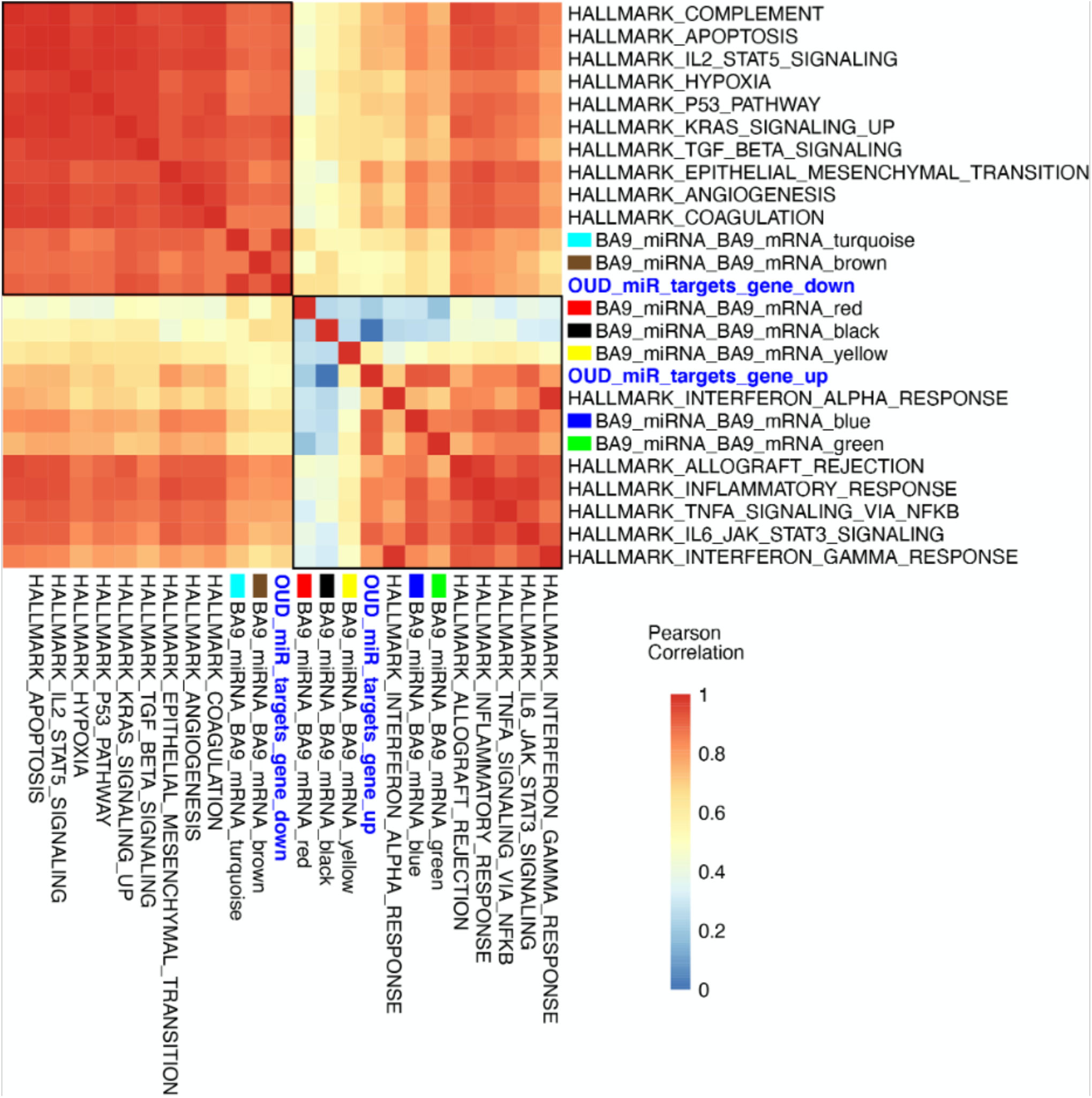
Gene set signatures for miRNA targets show robust and distinct correlation patterns associated with directionality. Gene signature scores were derived for OUD up-regulated and down-regulated miRNA targets, as well as for WGCNA modules, and comprehensive correlations were computed with the 50 Hallmark pathways over a collection of BA9 bulk tissue transcriptomes from 425 control brain transcriptomes from the GTEx project. Presented here is a subset of the Hallmark pathways, showing OUD down-regulated targets of miRNAs correlated with Hallmark pathways for hypoxia, angiogenesis, coagulation, and the TGF beta pathway, whereas OUD up-regulated targets of miRNAs correlated with immune system and inflammatory pathways.

## Discussion

In this study, we performed an integrated analysis of RNA-Seq and smallRNA-Seq data from same-subject postmortem brain (BA9) and blood tissues to explore the hypothesis that miRNAs and their mRNA targets are critical drivers of opioid-induced neurobiological alterations. Overall, we identified robust miRNA-mRNA networks in both BA9 brain and blood. Although there was no overlap between BA9 and blood differentially expressed miRNAs, we found strong overlap among the differentially expressed target genes of these miRNAs in both tissues. In addition, the corresponding enriched pathways derived from miRNA-mRNA networks in brain and blood were highly overlapping, with tube development, morphogenesis, and pathways related to endothelial cell function and vascular organization among the dominant enriched biological processes. Further, using WGCNA, we identified cell-type specific miRNA targets, particularly in astrocytes, neurons, and endothelial cells, associated with OUD transcriptomic dysregulation. The largest module identified by WGCNA (turquoise - comprised of 121 genes) was positively associated with endothelial cells, providing further evidence for a role of endothelial cells in opioid-induced brain alterations.

Of interest, the identified miRNA targets included genes we previously found to be involved in endothelial cell function, cytokine signaling, and angiogenesis pathways associated with OUD, including COL4A1, COL4A2, EGR1, EGR2, EGR4, FGF2, NR4A2, and DUSP10 (18). Among the differentially expressed and enriched miRNAs was 92a-3p, which has been previously found to be dysregulated in blood from male subjects after hydromorphone or oxycodone treatment (44). Our bioinformatics analysis identified miR-92a-3p to be significantly up-regulated in BA9, and its targets EGR1, EGR2, and DUSP10 (45, 46) to be significantly down-regulated. miR-92a is highly expressed in endothelial cells, which regulate vascular endothelial function. Overexpression of miR-92a in endothelial cells blocks angiogenesis and administration of a miR-92a inhibitor enhances recovery of damaged tissues in a mouse model of ischemia (47). Another identified miRNA of interest, miR-29b, was down-regulated in BA9, and its targets COL4A1 and COL4A2 were both up-regulated. In rodent model of cerebral infarction, miR-29 inhibited neuronal apoptosis (48). Disruption of miR-29 binding sites in COL4A1 lead to upregulation of COL4A1 and to ischemic cerebral small vessel disease (49).

Both miR-29b and miR-92a act through their targets via the p38 MAPK signaling pathway (50, 51), which we previously identified to be significantly enriched in OUD (18). Additional support for a role of the p38MAPK pathway in OUD pathophysiology was found by interrogation of the LINCS database with genes identified by our miRNA-mRNA networks in BA9 or blood. Although each network led to different top candidate drugs – TGFß receptor inhibitor for BA9 miRNAs, and p38 MAPK inhibitors for blood miRNAs – TGFß is a key upstream modulator of the p38 MAPK pathway (52). TGFß signaling can contribute to the pathogenesis of cardiovascular diseases (53), and TGFB2 was among the top differentially expressed genes in our previous study of gene dysregulation in OUD postmortem brain (41). Further, leveraging a collection of control brain transcriptomes from the GTEx project, we identified correlation of OUD miRNA targets with hypoxia, TGF beta, angiogenesis, coagulation, immune system, and inflammatory pathways.

Altogether, our current results support previous findings in human and animal studies of neurovascular alterations as a consequence of opioid abuse (54-64). Importantly, the American Heart Association recently advised on the risk of opioid use for neurovascular complications (65), underscoring the urgency of research aimed towards understanding mechanisms underlying opioid-induced neurobiological alterations. Further preclinical studies in animal or cell models are needed to clearly define these mechanisms.

In summary, our findings suggest that miRNA-driven mRNA dysregulation in OUD is profound and potentially realized through redundant and systemic alternatives. Our study demonstrates the utility of miRNAs to facilitate new avenues of mechanistic explorations that could lead to the development of novel therapeutic approaches to minimize or potentially reverse opioid-induced brain abnormalities.

## Acknowledgements

We are grateful for the invaluable donations and participation from families, as well as for the generous collaboration of the medical examiners at the Harris County Institute of Forensic Sciences. SLG, TG, RK, and CC were partially supported by The Cancer Prevention Institute of Texas (CPRIT) grants RP170005, RP210227, RP200504, NIH P30 shared resource grant CA125123, NIEHS grants P30 ES030285 and P42 ES027725, and NIMHD grant P50 MD015496. CWB was partially supported by grant R01DA044859.

---

**Supplemental Figure 1.**
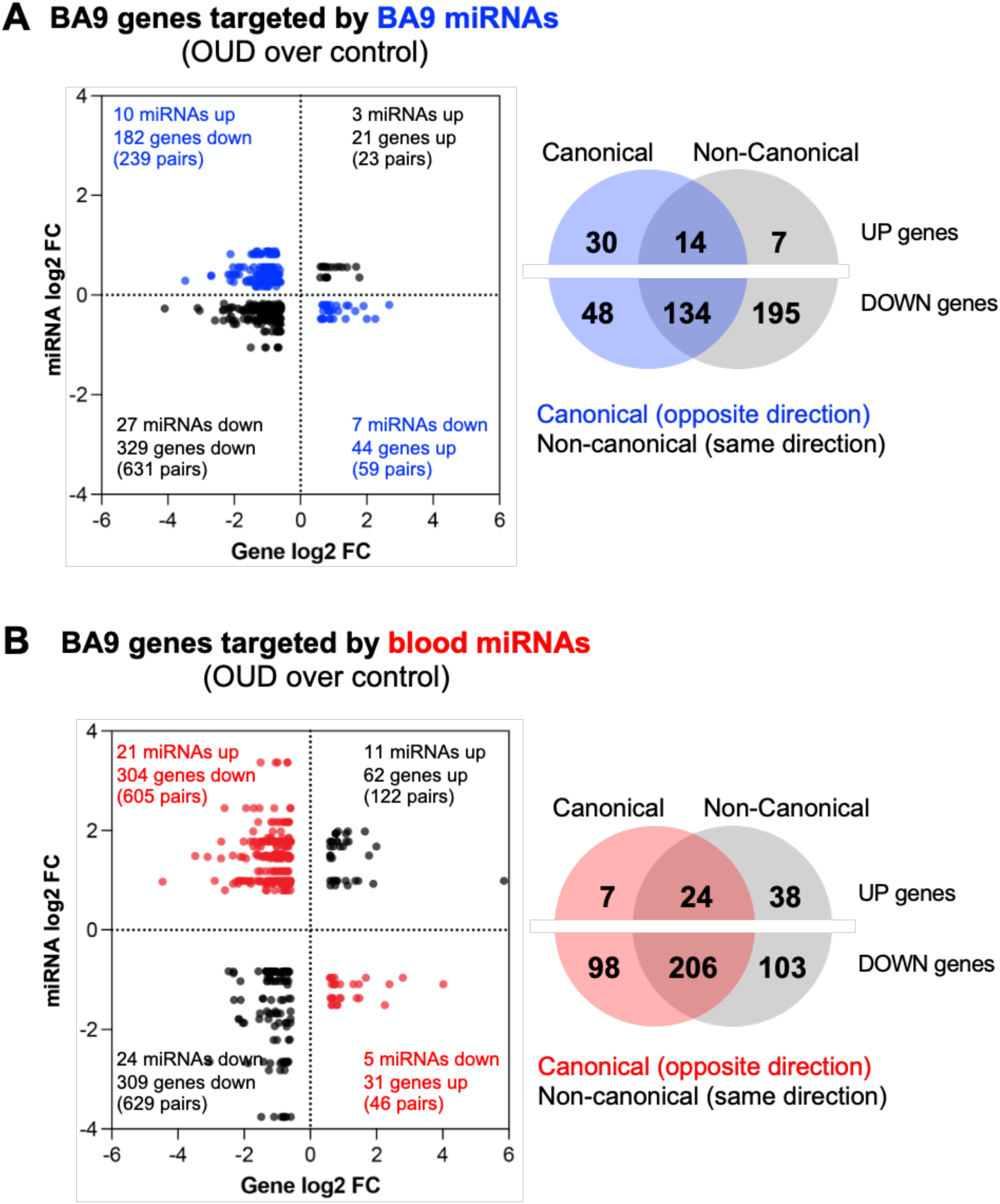
Evaluation of canonical and non-canonical miRNA-mRNA networks using BA9 and blood microRNAs. Using differentially expressed mRNAs measured in BA9, we assessed enrichment of miRNA targets based on differentially expressed miRNAs in (A) BA9 and (B) blood. We assessed canonical miRNA-mRNA regulation (e.g. opposite direction) and non-canonical miRNA-mRNA regulation (e.g. same direction). Some genes were enriched as miRNA targets under both canonical and non-canonical regulation.

**Supplemental Figure 2.**
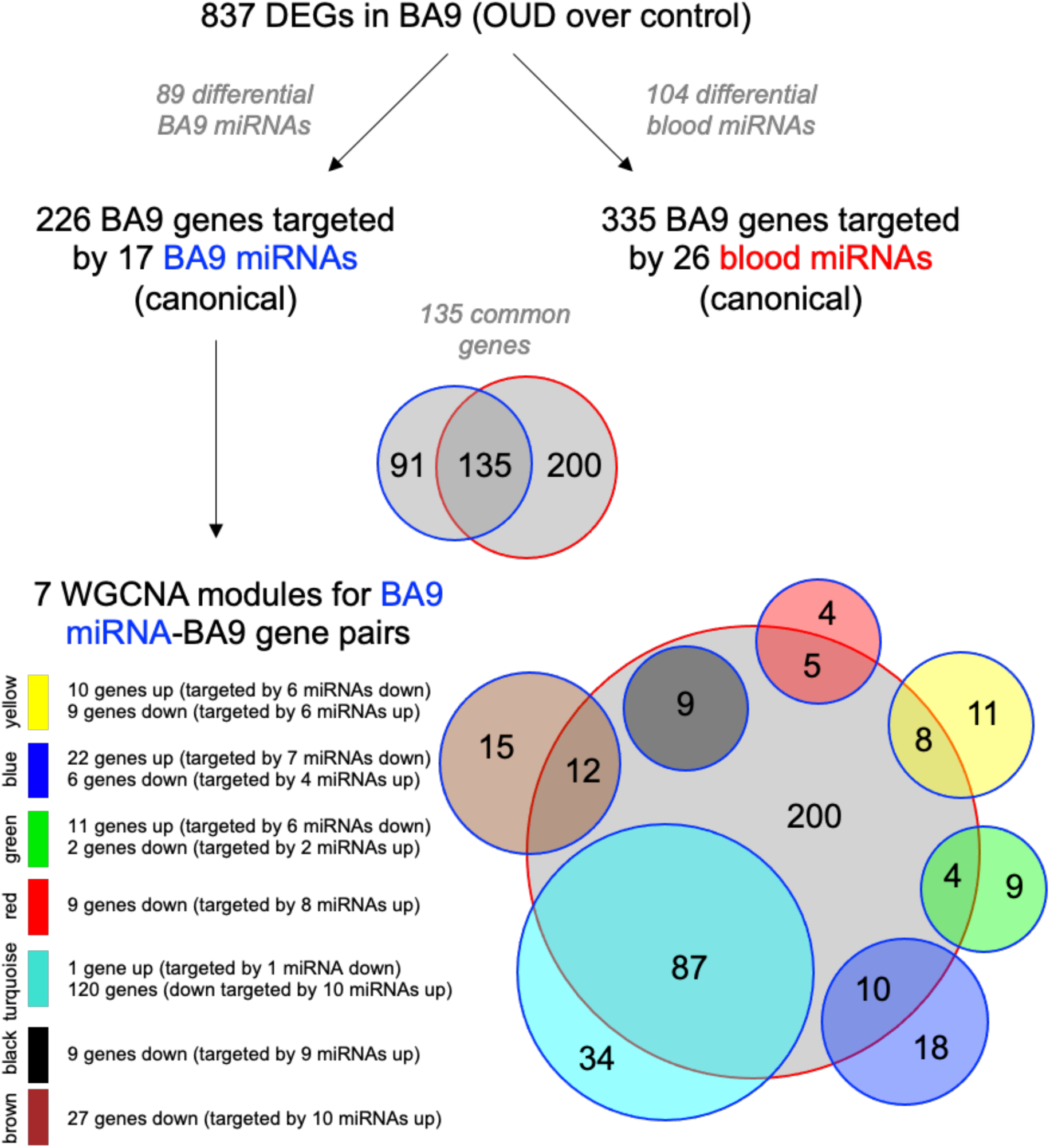
Overlap between WGCNA modules and targets of blood miRNAs. One hundred thirty-five gene targets were common between BA9 miRNAs and blood miRNAs. WGCNA BA9 module overlap with blood miRNA targets ranges between 31% (green module) to 100% (black module).

**Supplemental Figure 3.**
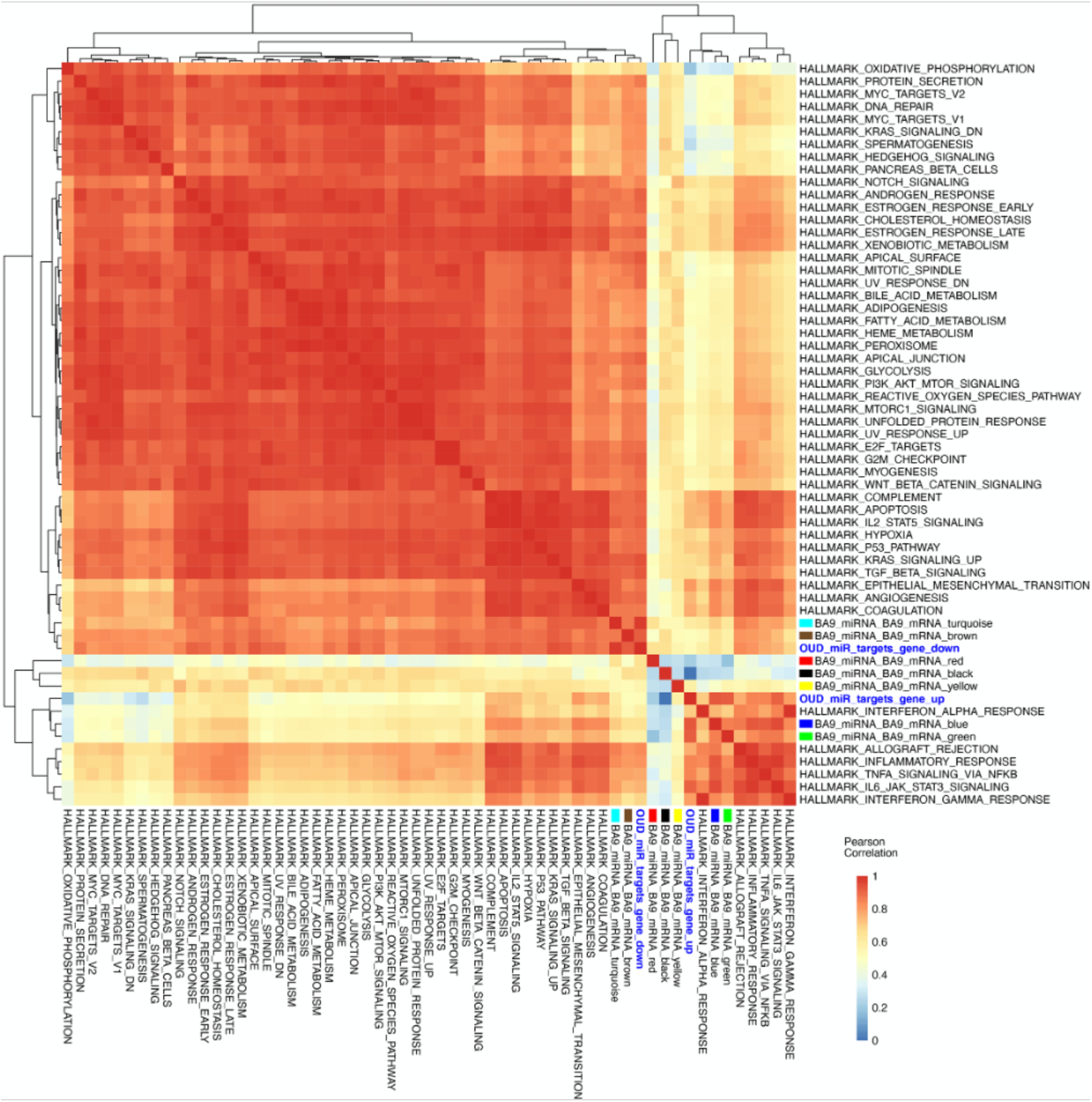
Comprehensive correlation map of gene set signatures for miRNA targets and Hallmark pathways. Gene signature scores were derived for OUD up-regulated and down-regulated miRNA targets, as well as for WGCNA modules, and comprehensive correlations were computed with the 50 Hallmark pathways over a collection of BA9 bulk tissue transcriptomes from 425 GTEx control brains.

## Supplemental Tables

**Supplemental Table 1. mRNA and miRNA network candidates. (**A) Coding genes significant at FDR<0.2, and fold change exceeding 1.5x. (B) Screening of candidate network miRNAs from BA9 at p<0.05. (C) Screening of candidate network miRNAs from blood at p<0.05.

**Supplemental Table 2. Genes targeted by miRNAs**. miRNA-mRNA pairs as determined using miRDB in BA9 brain (A) or blood (B).

**Supplemental Table 3. Pathway enrichment using genes targeted by miRNAs**. Enriched Gene Ontology Biological Process (GOBP) pathways by Over-Representation Analysis (ORA) using genes targeted by either BA9 or blood miRNAs. Significance (–log10 of the FDR), number of genes enriched for each pathway, and differentially expressed genes in each pathway are listed.

**Supplemental Table 4. WGCNA modules based on BA9 miRNA-mRNA network genes**. Membership of differentially expressed genes for each module is indicated. Overlap with blood miRNA-BA9 mRNA network genes, in either a canonical or non-canonical fashion, are annotated.

## Notes

### Competing Interest Statement

The authors have declared no competing interest.

